# A monoclonal antibody raised against human EZH2 cross-reacts with the RNA-binding protein SAFB

**DOI:** 10.1101/2023.04.03.535391

**Authors:** Rachel E. Cherney, Christine A. Mills, Laura E. Herring, Aki K. Braceros, J. Mauro Calabrese

## Abstract

The Polycomb Repressive Complex 2 (PRC2) is a conserved enzyme that tri-methylates Lysine 27 on Histone 3 (H3K27me3) to promote gene silencing. PRC2 is remarkably responsive to the expression of certain long noncoding RNAs (lncRNAs). In the most notable example, PRC2 is recruited to the X-chromosome shortly after expression of the lncRNA *Xist* begins during X-chromosome inactivation. However, the mechanisms by which lncRNAs recruit PRC2 to chromatin are not yet clear. We report that a broadly used rabbit monoclonal antibody raised against human EZH2, a catalytic subunit of PRC2, cross-reacts with an RNA-binding protein called Scaffold Attachment Factor B (SAFB) in mouse embryonic stem cells (ESCs) under buffer conditions that are commonly used for chromatin immunoprecipitation (ChIP). Knockout of EZH2 in ESCs demonstrated that the antibody is specific for EZH2 by western blot (no cross-reactivity). Likewise, comparison to previously published datasets confirmed that the antibody recovers PRC2-bound sites by ChIP-Seq. However, RNA-IP from formaldehyde-crosslinked ESCs using ChIP wash conditions recovers distinct peaks of RNA association that co-localize with peaks of SAFB and whose enrichment disappears upon knockout of SAFB but not EZH2. IP and mass spectrometry-based proteomics in wild-type and EZH2 knockout ESCs confirm that the EZH2 antibody recovers SAFB in an EZH2-independent manner. Our data highlight the importance of orthogonal assays when studying interactions between chromatin-modifying enzymes and RNA.

## Introduction

The Polycomb Repressive Complex 2 (PRC2) is a conserved histone-modifying enzyme that represses gene expression by catalyzing mono-, di-, and tri-methylation of Lysine-27 on Histone H3 (H3K27me3). PRC2 contains invariant subunits, including SUZ12, EED, and a catalytic engine − either EZH1 or EZH2 − as well as several auxiliary factors, which together serve to recruit PRC2 to specific genomic regions and modulate its catalytic activity. PRC2 is essential for mammalian development and is mutated in cancers and congenital disorders, underscoring its importance in gene regulation (Laugesen et al. 2016; Deevy and Bracken 2019; Yu et al. 2019).

PRC2 is highly expressed during early development, where it responds in a dramatic fashion to specific long noncoding RNAs (lncRNAs; (Schumacher et al. 1996; O’Carroll et al. 2001; Wang et al. 2001; Silva et al. 2003)). Within hours after the *Xist* lncRNA becomes expressed in the embryo proper (or in mouse embryonic stem cells; ESCs), PRC2 is recruited over the span of the 165 megabase (Mb) X chromosome (Plath et al. 2003; Okamoto et al. 2004; Zylicz et al. 2019). Expression of *Xist* transgenes from autosomal locations likewise result in chromosome-wide recruitment of PRC2, supporting the view that the *Xist* lncRNA itself, and not merely the act of its transcription, recruits PRC2 to chromatin (Trotman et al. 2021).

Exactly how PRC2 is recruited to chromatin by lncRNAs such as *Xist* remains unclear. While investigating this topic, we discovered that a widely used antibody raised against human EZH2 reproducibly cross-reacts with an RNA-binding protein called Scaffold Attachment Factor B (SAFB) in mouse ESCs during immunoprecipitations (IPs) but not by western blot. SAFB is a chromatin-associated RNA-binding protein that has been implicated in transcriptional repression and associates with the lncRNA *Xist* and may yet prove to play important roles in modulating PRC2 function (Townson et al. 2004; Mukhopadhyay et al. 2014; Chu et al. 2015; Bousard et al. 2019; Huo et al. 2020; McCarthy et al. 2021; Yu et al. 2021). Nevertheless, the data presented below highlight the importance of orthogonal assays in studying interactions between histone-modifying enzymes and RNA.

## Results and Discussion

By cross-linking immunoprecipitation (CLIP), PRC2 has been shown to associate with many nascent RNAs, a process that in certain cases may antagonize its enzymatic activity (Davidovich et al. 2013; Kaneko et al. 2013; Cifuentes-Rojas et al. 2014; Kaneko et al. 2014; Davidovich et al. 2015; Beltran et al. 2016; Wang et al. 2017; Beltran et al. 2019; Garland et al. 2019; Zhang et al. 2019; Long et al. 2020; Rosenberg et al. 2021). However, expression of the lncRNA *Xist* causes the near-immediate recruitment of PRC2 over the span of the inactive X-chromosome (Zylicz et al. 2019). Based on these data, we hypothesized that lncRNAs such as *Xist* recruit PRC2 indirectly, through RNA-binding proteins. Indeed, HNRNPK is an RNA-binding protein that appears to bridge PRC1 and *Xist*, providing precedent for a protein-bridging model for PRC2 (Pintacuda et al. 2017).

To determine whether PRC2 associates with RNA-binding proteins, we performed mass spectrometry-based proteomics (IP-MS) for EZH2 using a rabbit monoclonal antibody raised against human EZH2 from Cell Signaling Technology (CS5246). We prepared nuclear extracts from ESCs and performed IP-MS using EZH2-CS5246 under conditions in which the extracts were and were not pre-treated with RNase. Ethidium bromide staining of RNA prepared from these nuclear extracts confirmed degradation of RNA in the RNase treated samples (not shown). We identified all PRC2 major subunits, as well as many of its known accessory factors, and did not detect peptides for our most-enriched proteins in IgG control, which led us to infer that the antibody and our IP approach were successful (Figure 1A; Table S1). Notably, IP with EZH2-CS5246 identified several RNA-binding proteins under both RNase-free and RNase-treated conditions. The most highly-ranked of these was SAFB, which has previously been found to interact with EZH2 in human and mouse cells (Gao et al. 2012; Cao et al. 2014; Oksuz et al. 2018) and has been proposed to be necessary for H3K27me3 deposition at certain androgen-repressed genes (Mukhopadhyay et al. 2014). SAFB has also been shown to interact with *Xist* in mouse and human ESCs (Chu et al. 2015; Bousard et al. 2019; Yu et al. 2021). Using CS5246, we verified the interaction between EZH2 and SAFB via IP-western (Figure 1B). EZH2-CS5246 IP retrieved SAFB, yet SAFB IP retrieved trace amounts of EZH2; likewise, IP with another core component of PRC2, SUZ12, did not retrieve SAFB (Figure 1B). Nevertheless, the prior connections between SAFB, EZH2, and *Xist* were intriguing enough that we elected to study the interaction further.

**Figure 1.**
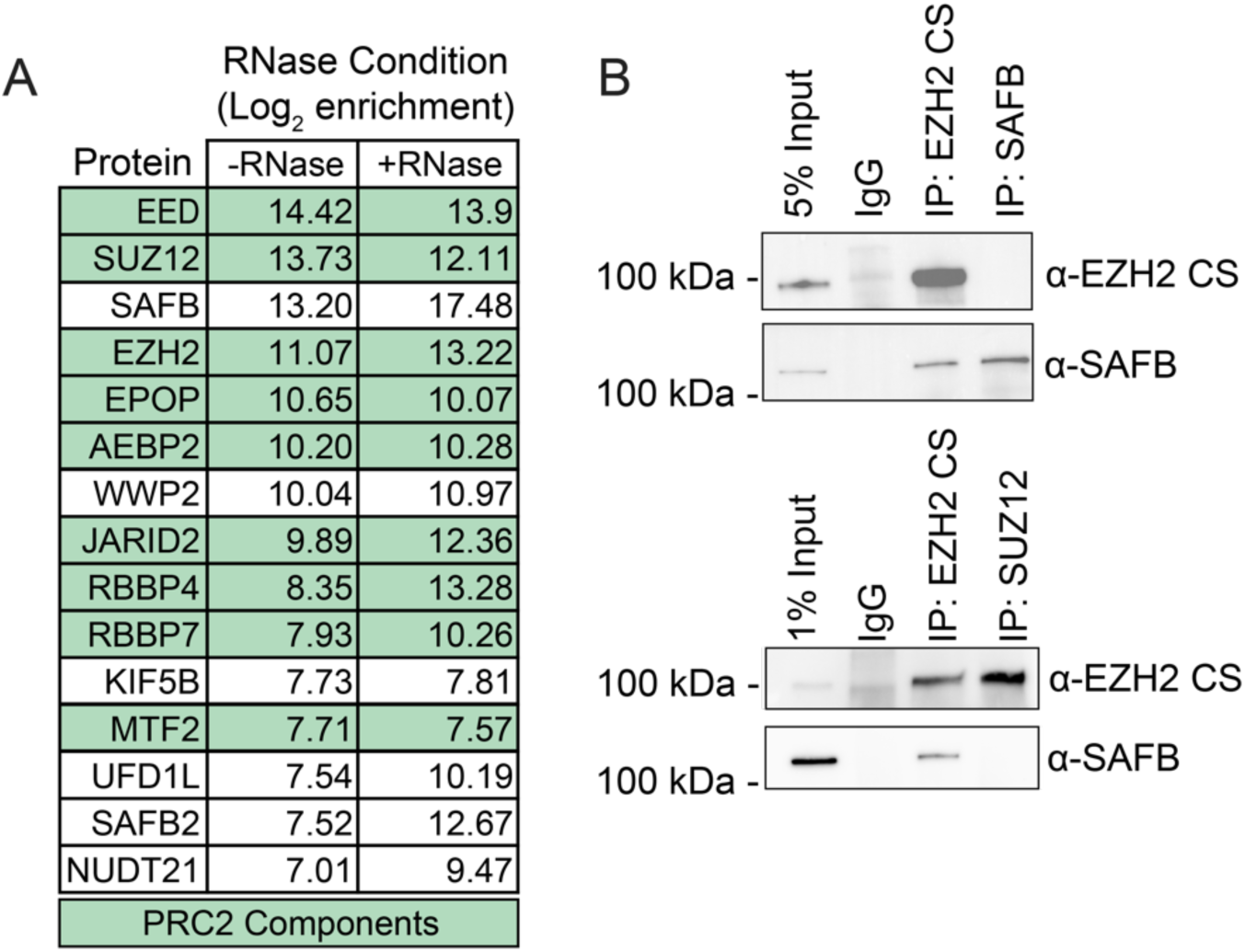
IP-MS with EZH2-CS5246 identifies PRC2 complex members and SAFB. **(A)** Log2 enrichment of top 15 proteins identified via IP-MS with EZH2-CS5246 in samples treated with or without RNase. PRC2 components are highlighted in green. IP-MS data can be found in Table S1. **(B)** IP-western analysis using EZH2-CS5246, SAFB, and SUZ12 antibodies.

While CLIP data show that PRC2 directly binds RNA somewhat promiscuously *in vivo*, we reasoned that specific, bridged interactions between RNA, RNA-binding proteins, and PRC2 could be missed by CLIP, which was designed to detect direct RNA/protein interactions. In contrast, bridged interactions might be apparent in RNA-IP (RIPs) from formaldehyde-crosslinked cells, which, in addition to direct interactions, can also reveal indirect ones in the form of RNA bound to proteins bound to the protein-target of the IP (Hoffman et al. 2015). To identify the RNAs retrieved by EZH2-CS5246 *in vivo*, we used a formaldehyde-based RIP-Seq protocol (Schertzer et al. 2019a), which relies on a series of washes that are also used in many chromatin immunoprecipitation (ChIP) protocols, including: one five-minute wash in a buffer that contains 500mM NaCl, 1% Triton X-100, 0.1% sodium deoxycholate, and 0.1% SDS; and another five-minute wash in a buffer that contains 250mM LiCl, 0.5% sodium deoxycholate, and 0.5% NP-40. Both wash conditions are predicted to destabilize non-specific protein-protein interactions. In parallel, we used the same RIP protocol to retrieve RNA associated with an endogenous SAFB antibody and mouse IgG as a negative control. By RIP-Seq, we observed a striking co-localization between the RNA retrieved EZH2-CS5246 and SAFB. Thirty-two percent of all EZH2 RIP peaks overlapped with SAFB RIP peaks (10,813/33,671 EZH2 peaks; p<2e-16 hypergeometric test; Figure 2A; Table S2). When examining the top 10,000 EZH2 RIP peaks (ranked by EZH2 RIP signal), the overlap increased to eighty percent (7,968/10,000 EZH2 peaks; p<2e-16 hypergeometric test). Concordantly, SAFB and EZH2 RIP signal under EZH2 RIP peaks were highly correlated (Figure 2B). Because transient depletion of SAFB by inducible CRISPR resulted in loss of EZH2-CS5246 signal at a subset of target lncRNAs (Figure 2C), we elected to knockout SAFB and its paralogue SAFB2 in ESCs. Generation and study of SAFB/2 double-knockout (DKO) ESCs are described in greater depth in (Cherney et al. 2022). We performed RIP-Seq using EZH2-CS5246 in wild-type and SAFB/2 DKO ESCs and observed a loss of signal at essentially all SAFB- and EZH2-CS5246-enriched regions in SAFB/2 DKO ESCs, indicating that RIP with EZH2-CS5246 retrieves RNA in a SAFB-dependent manner (Figure 2D-F).

**Figure 2.**
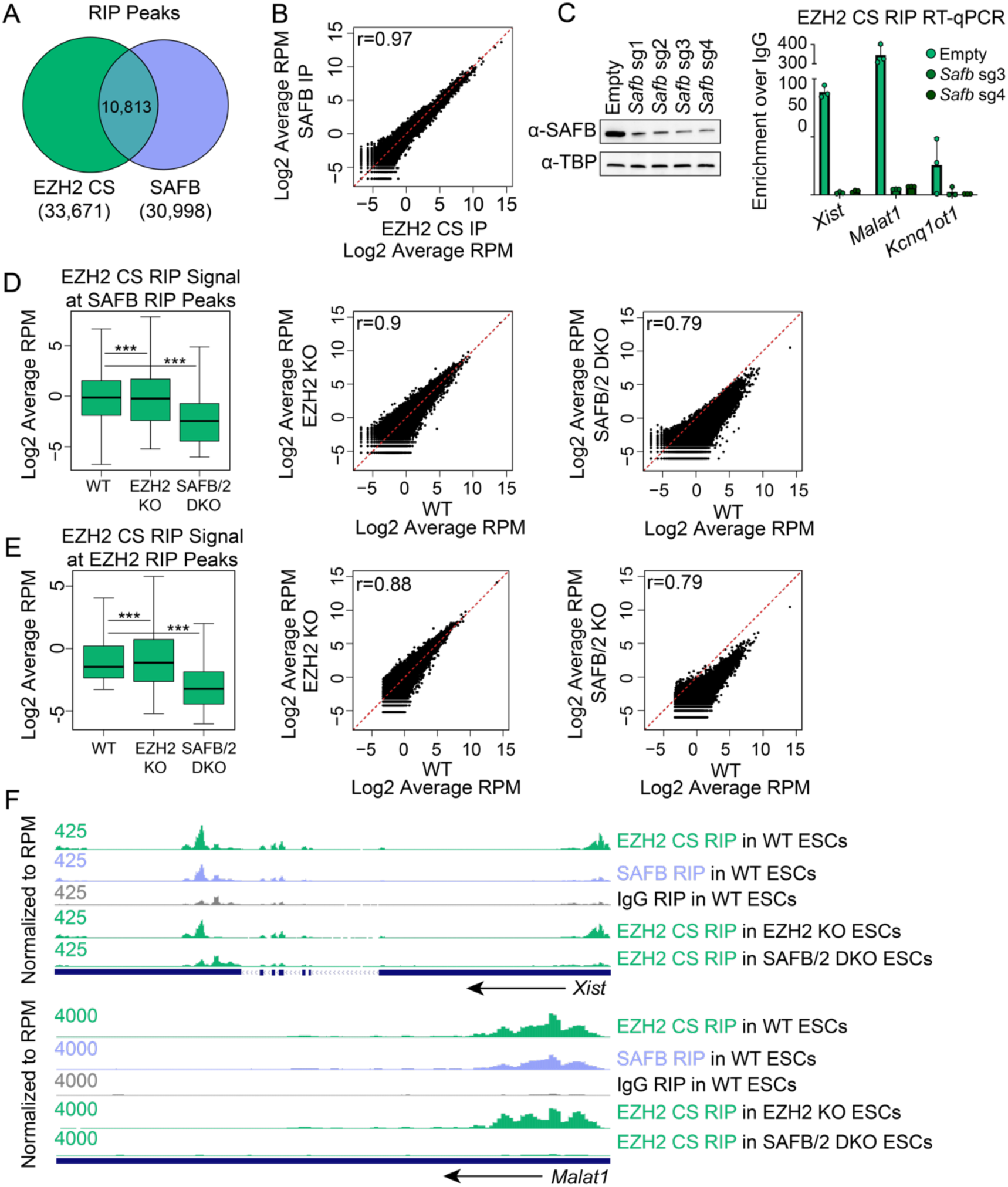
EZH2-CS5246 recovers SAFB-dependent signal in RIP. **(A)** Locational overlap between EZH2-CS5246 and SAFB RIP peaks. Hypergeometric distribution p-value, <2e-16. EZH2 RIP peaks can be found in Table S2. **(B)** SAFB and EZH2-CS5246 RIP signal under EZH2-CS5246 RIP peaks. Pearson’s r=0.97. **(C)** Western blot showing SAFB levels following three days of Cas9 induction. RT-qPCR detecting *Xist*, *Malat1*, and *Kcnq1ot1* in EZH2-CS5246 RIP from ESCs stably expressing a doxycycline-inducible Cas9 and no sgRNA (Empty) or one of two separate sgRNAs targeting SAFB. **(D)** EZH2-CS5246 RIP signal under SAFB RIP peaks in WT, EZH2 KO and SAFB/2 DKO cell lines. p-value, *** = <2e-16, t-test. **(E)** EZH2-CS5246 RIP signal under EZH2 RIP peaks in WT, EZH2 KO and SAFB/2 DKO cell lines. p-value, *** = <2e-16, t-test. **(F)** UCSC wiggle tracks of EZH2 and SAFB RIP signal in WT, EZH2 KO and SAFB/2 DKO ESCs over the lncRNAs *Xist* and *Malat1*.

However, we remained concerned that previously, we did not retrieve SAFB in SUZ12 IPs, nor did we reproducibly retrieve EZH2 by SAFB IP (Figure 1B). For this reason, we used CRISPR to generate EZH2 knockout (KO) ESCs. By PCR, RT-PCR, and western blot with EZH2-CS5246 we observed complete loss of EZH2 DNA, RNA, and EZH2 protein in EZH2 KO ESCs and no evidence of cross-reactivity with other proteins in the EZH2-CS5246 western blot (Figure 3A-C). Nonetheless, RIP-Seq using EZH2-CS5246 showed essentially no change in signal at EZH2-CS5246-enriched regions in wild-type versus EZH2 KO ESCs (Figure 2E and 2F; Table S2). Moreover, by IP-western and IP-MS, EZH2-CS5246 still retrieved SAFB even in EZH2 KO ESCs (Figure 3D-F; Table S1). We conclude that under our RIP conditions, but not in western blots, EZH2-CS5246 cross-reacts with SAFB, and that by RIP, the RNA retrieved by EZH2-CS5246 requires the presence of SAFB and not EZH2.

**Figure 3.**
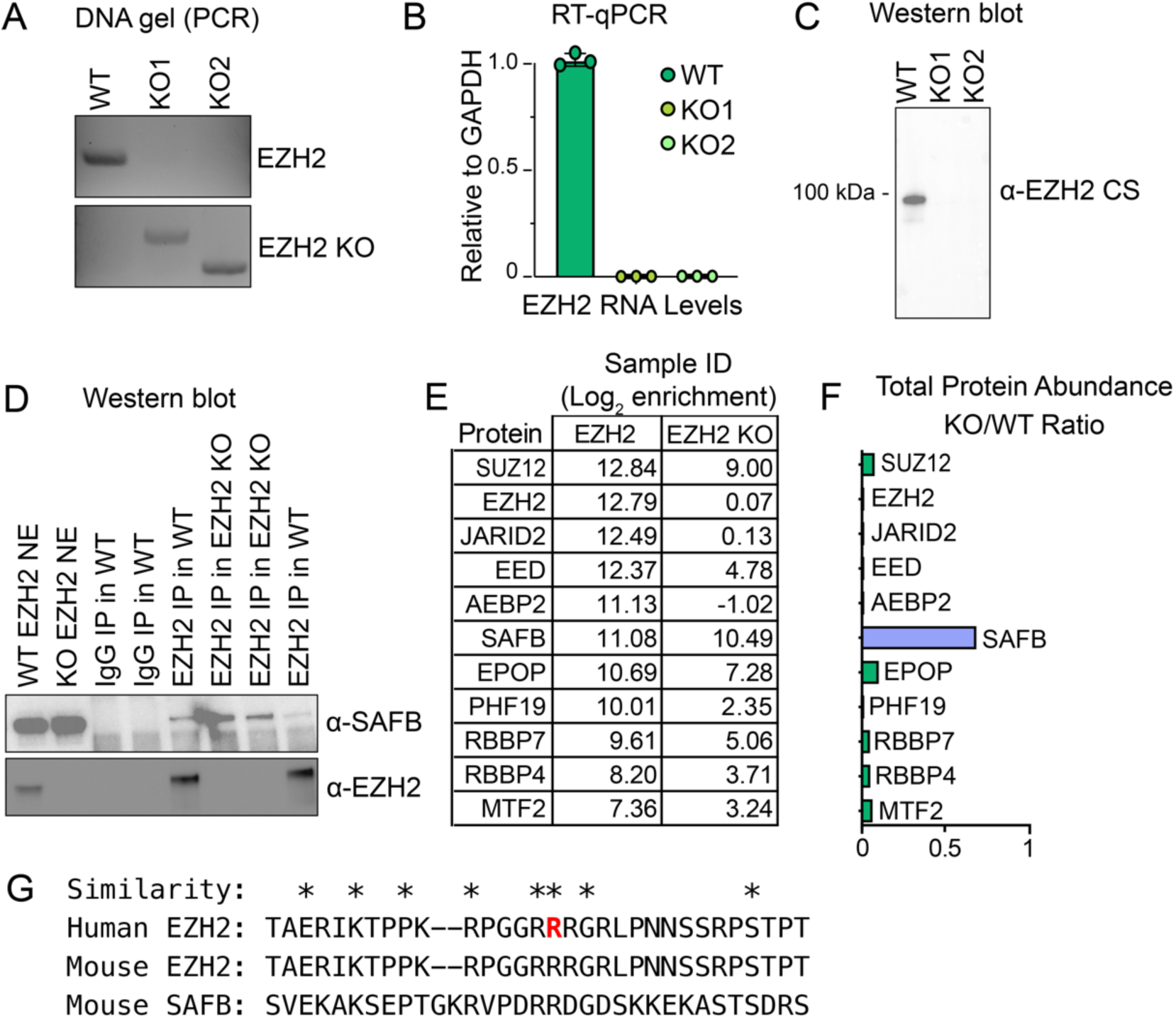
EZH2-CS5246 pulls down SAFB in EZH2 KO ESCs. **(A)** PCR validation of EZH2 KO. **(B)** RT-qPCR of EZH2 RNA levels in WT and two independent EZH2 KO clones. **(C)** Western blot using EZH2-CS5246 in WT and two independent EZH2 KO clones. **(D)** IP-western from WT and EZH2 KO nuclear extracts using EZH2-CS5246 or IgG control. **(E)** Log2 enrichment of PRC2 members and SAFB in EZH2-CS5246 IP-MS from WT and EZH2 KO nuclear extracts. IP-MS protein lists and quantification can be found in Table S1. **(F)** Ratio of protein abundance between WT and EZH2 KO nuclear extracts of PRC2 members and SAFB. **(G)** Sequence alignment of human EZH2-CS5246 epitope region, the analogous region from mouse EZH2, and a predicted region of similarity in mouse SAFB. The EZH2-CS5246 epitope surrounds Arg354 of human EZH2 and is denoted in red.

We extracted the approximate epitope used to generate EZH2-CS5246 and aligned it to SAFB and observed no significant linear sequence similarity between the epitope and SAFB. However, we note that the approximate EZH2-CS5246 epitope does harbor several arginine and glycine residues, and regions of SAFB are also enriched in R/G residues, providing a possible reason for the cross-reactivity of EZH2-CS5246 with SAFB (Figure 3G).

Because EZH2-CS5246 has been used in hundreds of publications (632 at the time of this writing), we wanted to verify that the antibody retrieves EZH2-dependent signal by ChIP-Seq. We note that the wash conditions we used for RIP-Seq in this study are either identical or similar to wash conditions previously used for EZH2-CS5246 ChIP-Seq in our own and other works (Deaton and Bird 2011; Kaneko et al. 2013; Mu et al. 2018; Schertzer et al. 2019a). We performed ChIP-Seq with EZH2-CS5246 in ESCs and compared our data to two previous studies in ESCs, one using EZH2-CS5246 (Mu et al. 2018) and another using an EZH2 antibody generated in-house (Kaneko et al. 2013)(Table S5). The three datasets were highly concordant by eye (Figure 4A), identified near-complete overlap in peak locations (Figure 4B), and exhibited highly significant correlations when examining signal under peaks (Figure 4C), supporting the view that EZH2-CS5246 can be used to generate PRC2-specific signal in ChIP assays.

**Figure 4.**
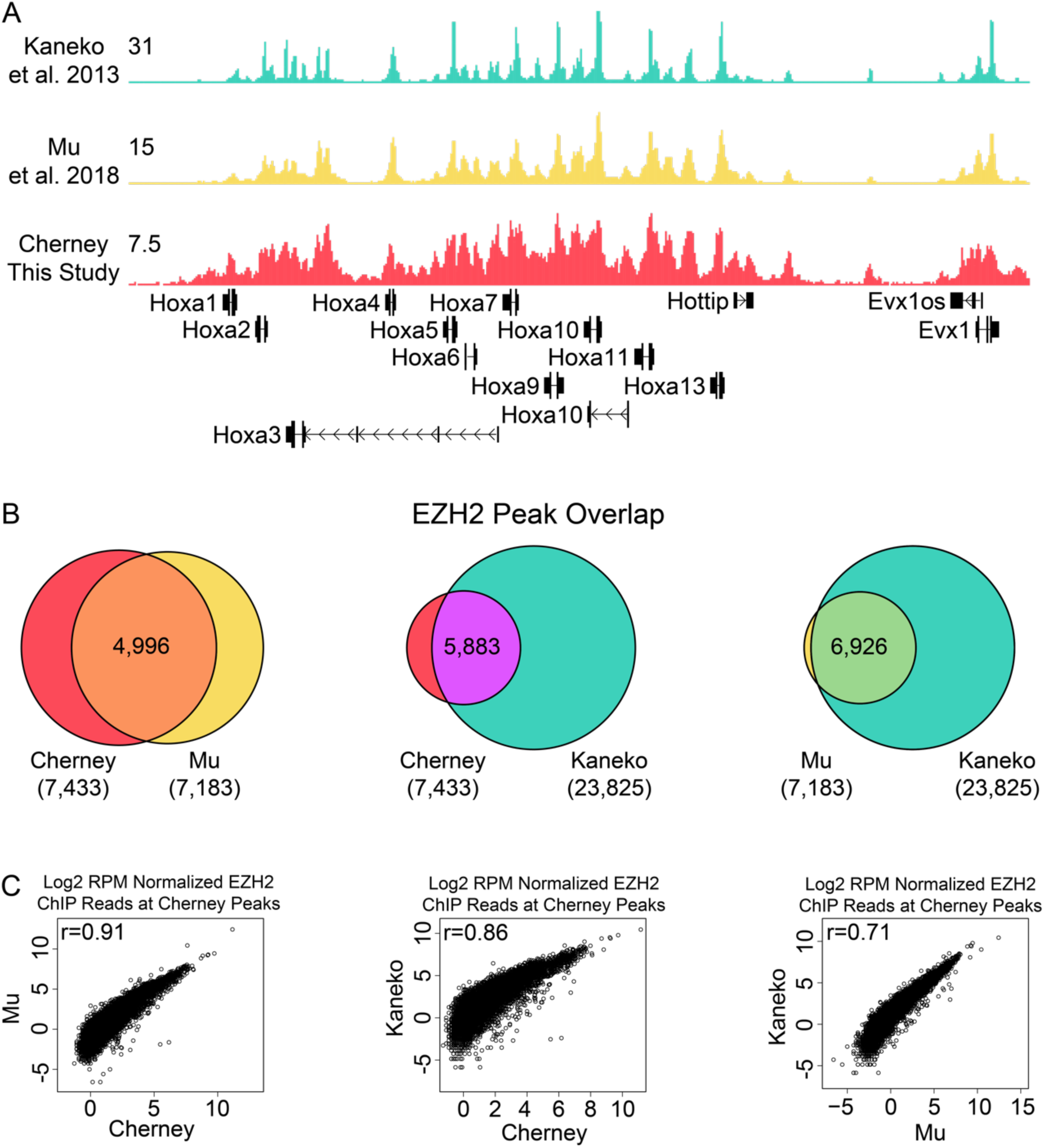
Correlations between EZH2 ChIP-Seq data sets. **(A)** EZH2 ChIP-Seq UCSC wiggle tracks of HOXA cluster for three EZH2 ChIP data sets. (Kaneko et al. 2013) used an EZH2 antibody generated in-house; (Mu et al. 2018) and Cherney (this study) used EZH2-CS5246. **(B)** Peak overlap and **(C)** Correlation of signal under Cherney peaks between EZH2 ChIP-Seq data sets. Cherney peak locations are listed in Table S5.

PRC2 exhibits an enigmatic relationship with RNA (Trotman et al. 2021). By performing IP with a rabbit monoclonal antibody that is highly specific for EZH2 by western blot and yields specific signal for PRC2 by ChIP, we retrieved SAFB, a chromatin-associated RNA-binding protein that has previously been implicated in gene silencing and PRC2 function, and which has been previously shown to associate with *Xist/XIST* (Townson et al. 2004; Mukhopadhyay et al. 2014; Chu et al. 2015; Bousard et al. 2019; Huo et al. 2020; McCarthy et al. 2021; Yu et al. 2021). We find that the RNA enriched in EZH2-CS5246 RIPs is SAFB- and not EZH2-dependent. SAFB or other RNA-binding proteins may yet prove to be important for RNA-mediated recruitment of PRC2 to chromatin. Indeed, RBFOX2 and HNRNPA2B1 may be two such proteins (Meredith et al. 2016; Wei et al. 2016). However, continued work in this area is needed to arrive at mechanistic clarity. Our study highlights the importance of orthogonal assays when interpreting interactions between RNA and chromatin-modifying enzymes, including and not limited to PRC2.

## Methods

### Experimental methods

#### Cell culture

Male mouse ESCs expressing doxycycline-inducible *Xist* from the *Rosa26* locus (derivation described in (Trotman et al. 2020)) were grown in DMEM (Gibco) supplemented with 15% Qualified Fetal Bovine Serum (Gibco), 1% Pen/Strep (Gibco), 1% L-Glutamine (Gibco), 1% Non-Essential Amino Acids (Gibco), 100μM betamercaptoethanol (Sigma) and 0.2% LIF. Cells were passaged every other day and maintained in incubators set at 37°C and 5% CO_2_. Media was replaced daily.

#### Nuclear Extraction of ESCs

ESC nuclear extracts were prepared as in (Carey et al. 2009). All nuclear extraction steps were performed at 4°C. Briefly, cells were grown to ∼80% confluency in 15cm plates and washed twice in cold 1xPBS. 10mL of 1xPBS supplemented with 0.5mM PMSF (Thermo Fisher #36978) per plate were scraped and consolidated to a 15mL conical tube. Cells were spun at 1000 rpm for 5 minutes. PBS was aspirated and cells were resuspended in 2x Packed Cell Volume of hypotonic cell lysis buffer (10mM HEPES-KOH pH 7.9, 1.5mM MgCl_2_, 10mM KCl) supplemented with 0.5mM DTT (ThermoFisher #15508013), 0.5mM PMSF (Thermo Fisher #36978) and 1x Protease Inhibitor Cocktail (PIC; Sigma Product #P8340); ex: 100μL of pelleted cells would be resuspended in 200μL hypotonic cell lysis buffer with supplements. NP-40 was added to the cell resuspension to a final concentration of 0.1% and pellet was incubated for 5 min. Resuspended cell pellet was transferred to a B-Dounce and was lysed via Dounce homogenizer ten times, after which the lysed cells were spun down for 4 minutes at 2,000 rpm and supernatant removed. Cells were then resuspended in 1.5x Packed Cell Volume of nuclear lysis buffer (20mM HEPES-KOH pH 7.9, 25% Glycerol, 420mM KCl, 1.5mM MgCl_2_, 0.2mM EDTA) supplemented with 0.5mM DTT (ThermoFisher #15508013), 0.5mM PMSF (Thermo Fisher #36978) and 1x Protease Inhibitor Cocktail (PIC; Sigma Product #P8340) and incubated for 30 minutes with rotation, then spun down for 5 minutes at 5,000 rpm. Supernatant was moved to a new tube and extract was fully cleared by spinning at top speed for 15 minutes. Nuclear extraction was repeated by pellet resuspension in 1.5x Packed Cell Volume of nuclear lysis buffer supplemented with 0.5mM DTT (ThermoFisher #15508013), 0.5mM PMSF (Thermo Fisher #36978) and 1x Protease Inhibitor Cocktail (PIC; Sigma Product #P8340) and incubated for 30 minutes with rotation, then spun down for 5 minutes at 5,000 rpm. Supernatant was moved to a new tube and extract was cleared by spinning at top speed for 15 minutes. Nuclear extract was then dialyzed for two hours in Dialysis Buffer (20mM HEPES-KOH pH 7.9, 20% Glycerol, 100mM KCl, 0.2mM EDTA} supplemented fresh 0.2mM DTT, on a rotating platform at 4°C using a Mini-Dialyzer (Thermo Fisher #88403). After two hours, dialysis buffer with 0.2mM DTT was replaced, and samples were dialyzed overnight on a rotating platform at 4°C. The next day supernatant was cleared by spinning down at high speed for 15 minutes. Samples were snap frozen in 200μL aliquots in liquid N_2_ and stored at -80°C until use.

#### IP-MS sample preparation

40μL protein A/G agarose beads (Santa Cruz sc-2003) were washed 3 times in blocking buffer (0.5% BSA in 1xPBS) and incubated overnight at 4°C with 20μL EZH2 antibody (anti-EZH2, Cell Signaling #5246) or 20μg IgG (anti-IgG; Invitrogen #02-6102). Beads were then washed 3 times in fRIP buffer (25mM Tris-HCl pH 7.5, 5mM EDTA, 0.5% NP-40, 150mM KCl). To the beads, 3.2mg of WT ESC or EZH2 KO ESC nuclear extract was added. RNase treated nuclear extracts were incubated with 1μL:40μL ratio of RNase (Thermo Fisher EN0531) to nuclear extract for one hour at 4°C with rotation before being added to the antibody conjugated beads. Physiological salt levels (175mM) were recapitulated by bringing nuclear extracts to a total of 2.5mL in 1:1 RIPA (50mM Tris-HCl, pH8.0, 1% Triton X-100, 0.5% sodium deoxycholate, 0.1% SDS, 5mM EDTA, 150mM KCl) and fRIP buffers (25mM Tris-HCl pH 7.5, 5mM EDTA, 0.5% NP-40, 150mM KCl), supplemented with 1x PIC (PIC; Sigma Product #P8340), 12.5μL SuperaseIN (Thermo Fisher #AM2696), and 0.5mM DTT (ThermoFisher #15508013). Samples were rotated overnight at 4°C and then washed once with 1mL fRIP buffer, resuspended in 1mL PolII ChIP Buffer (50mM Tris-HCl pH 7.5, 140mM NaCl, 1mM EDTA, 1mM EGTA, 1% Triton X-100, 0.1% Na-deoxycholate, 0.1% SDS), and transferred to a new tube. Samples were then rotated for 5 minutes and spun down at 2,000 rpm for one minute. Samples were washed two more times with PolII ChIP Buffer, once with High Salt ChIP Buffer (50mM Tris-HCl pH7.5, 500mM NaCl, 1mM EDTA, 1mM EGTA, 1% Triton X-100, 0.1% Na-deoxycholate, 0.1% SDS), and resuspended in 1mL LiCl buffer (20mM Tris pH 8.0, 1mM EDTA, 250mM LiCl, 0.5% NP-40, 0.5% Na-deoxycholate), with a 5 minute rotation and 1 minute spin down at 2,000 rpm after each wash. Samples were moved to a new tube at the final LiCl wash. After aspiration of the LiCl buffer, samples were resuspended in cold 1xPBS and moved to a new tube. Samples were washed 3 times with 1mL cold 1xPBS. 25% of samples were saved for western blot, and the remaining 75% were subjected to on-bead trypsin digestion. (Rank et al. 2021). Briefly, after the last wash buffer step during affinity purification, beads were resuspended in 50μL of 50mM ammonium bicarbonate (pH8). On-bead digestion was performed by adding 50μL 50mM ammonium bicarbonate (pH8) and 1μg trypsin and incubated, shaking, overnight at 37°C. Beads were pelleted and transferred supernatants to fresh tubes. The beads were washed twice with 100μL LC-MS grade water, and washes added to the original supernatants. Samples were acidified by adding formic acid to final concentration of 2%, to pH ∼2. Peptides were desalted using peptide desalting spin columns (Thermo), lyophilized, and stored at -80°C until further analysis.

#### LC/MS/MS analysis

The peptide samples were analyzed by LC/MS/MS using an Easy nLC 1200 coupled to a QExactive HF Biopharma mass spectrometer (Thermo Scientific). Samples were injected onto an Easy Spray PepMap C18 column (75μm id × 25cm, 2μm particle size) (Thermo Scientific) and separated over a 2 hr method. The gradient for separation consisted of 5–45% mobile phase B at a 250 nl/min flow rate, where mobile phase A was 0.1% formic acid in water and mobile phase B consisted of 0.1% formic acid in acetonitrile (ACN). The QExactive HF was operated in data-dependent mode where the 15 most intense precursors were selected for subsequent fragmentation. Resolution for the precursor scan (m/z 350–1700) was set to 60,000, while MS/MS scans resolution was set to 15,000. The normalized collision energy was set to 27% for HCD. Peptide match was set to preferred, and precursors with unknown charge or a charge state of 1 and ≥ 7 were excluded.

#### Immunoprecipitation followed by western blot (IP-western)

25μL protein A/G agarose beads (Santa Cruz sc-2003) were washed 3 times in blocking buffer (0.5% BSA in 1xPBS) and incubated overnight at 4°C with 7.5μL antibody (anti-EZH2, Cell Signaling #5246; SAFB, Bethyl A300-812A; SUZ12, Cell Signaling #3737) or 3μL (7.5μg) IgG (anti-IgG; Invitrogen #02-6502). The following day, beads were washed 2x in blocking buffer and resuspended in 300μg nuclear extract. Nuclear extract was diluted with a 1:1 mixture of RIPA (50mM Tris-HCl, pH8.0, 1% Triton X-100, 0.5% sodium deoxycholate, 0.1% SDS, 5mM EDTA, 150mM KCl) and fRIP buffers (25mM Tris-HCl pH 7.5, 5mM EDTA, 0.5% NP-40, 150mM KCl) to bring salt concentrations to physiological levels (175mM). Nuclear extract mixture was supplemented with 1mM PMSF (Thermo Fisher #36978) and 1x Protease Inhibitor Cocktail (PIC; Sigma Product #P8340) and samples rotated overnight at 4°C. The next day, samples were washed four times in a 1:1 mixture of RIPA (50mM Tris-HCl, pH8.0, 1% Triton X-100, 0.5% sodium deoxycholate, 0.1% SDS, 5mM EDTA, 150mM KCl) and fRIP buffers (25mM Tris-HCl pH 7.5, 5mM EDTA, 0.5% NP-40, 150mM KCl) to bring salt concentrations to physiological levels (175mM). Nuclear extract mixture was supplemented with 1mM PMSF (Thermo Fisher #36978) and 1x Protease Inhibitor Cocktail (PIC; Sigma Product #P8340). Tubes were changed for the first and the last wash. After the last wash, beads were spun down and resuspended in 75μL 4xSDS loading buffer (Sigma Aldrich Recipe: 0.2M Tris-HCl pH6.8, 0.4M DTT, 8% (w/v) SDS, 6mM Bromophenol blue, 4.3M Glycerol) diluted to 1x, and 20μL loaded for western blotting.

#### Whole-cell western blots

To isolate protein for western blotting, 0.8x10e6 cells were washed with 1xPBS and then lysed with 500μL RIPA buffer (10mM Tris-Cl pH7.5, 1mM EDTA, 0.5mM EGTA, 1% NP40, 0.1% sodium deoxycholate, 0.1% Sodium Dodecyl Sulfate, 140mM Sodium Chloride) supplemented with 1mM PMSF (Thermo Fisher #36978) and 1x Protease Inhibitor Cocktail (PIC; Sigma Product #P8340). Cell suspensions were rotated for 15 minutes at 4°C, then spun down at high speed at 4°C for 15 minutes and supernatant was collected. Prior to western blotting, protein levels were quantified using the DC assay from Biorad (Product #5000006). 4x SDS loading buffer (Sigma Aldrich Recipe: 0.2M Tris-HCl pH6.8, 0.4M DTT, 8% (w/v) SDS, 6mM Bromophenol blue, 4.3M Glycerol) was added to samples to 1x final concentration. Samples were then boiled for 5 minutes at 95°C, and equal μg (whole cell lysate) or μL (immunoprecipitation) amounts were loaded onto BioRad TGX Stain Free Gels. Samples were run at 50V until past stacking gel, then at 150V for 1-2 hrs. Gels were transferred to PVDF (Immobulon #IPVH00010) membrane either for 1 hour at 125V at 4°C or overnight at 25V at 4°C. Membranes were blocked for 45 minutes in 1xTBST + 5% milk (Biorad #1706404). Membranes were then incubated with primary antibody either overnight at 4°C or for 1-3 hours at RT. Membranes were washed 3x for 5 minutes each in 1x TBST. Secondary antibodies were diluted in 1xTBST + 5% milk and incubated with membranes for 45 minutes (1:100,000; Invitrogen). Membranes were then washed 3x in 1xTBST washes for 10 minutes, before being imaged with ECL (Thermo Fisher #34096). Antibodies used were: EZH2 (Cell Signaling #5246, 1:1000), EZH2 (Cell Signaling #3147, 1:1000), SAFB (Bethyl A300-812A, 1:3000), TBP (Abcam ab818, 1:2000), Goat anti-Mouse (Thermo Scientific A16072, 1:100,000) and Goat anti-Rabbit (Thermo Scientific G21234, 1:100,000).

#### Inducible knockdown of SAFB in ESCs

Guide RNAs targeting SAFB were designed using Benchling. sgRNA sequences are found in Table S3. Guides were cloned into the rtTA-BsmbI piggyBac vector from ((Schertzer et al. 2019b); Addgene plasmid #126028). For stable plasmid integration, parent ESCs were seeded at 0.5x10^6^ cells per well in a 6-well plate. The following day, the cells were transfected using Lipofectamine 3000 (Invitrogen L3000-015), using the following amounts of reagent: 1818ng of sgRNA or empty plasmid, 455ng Cas9 plasmid, and 228ng Transposase (2.5μg total), mixed with 5μL P3000 reagent, 7.5μL Lipofectamine 3000 reagent, and with Opti-MEM media (Gibco #31985-070) to final volume of 250μL. The reagents were incubated for 5 minutes at room temperature before being added to cells with fresh media. After 24 hours, cells were co-selected with Hygromycin (150μg/mL) for one week and G418 (200μg/mL) for 12 days. To deplete SAFB from cells, fresh doxycycline was added to media daily at 1μg/mL for three days. Media was replaced daily.

#### Generation of EZH2 knockout ESCs

sgRNAs to delete the EZH2 locus were designed to the mm10 genome using CRISPOR with the specifications: 20bp-NGG – Sp Cas9, Sp Cas9-Hf1, eSp Cas9 1.1 (Concordet and Haeussler 2018). sgRNA sequences are found in Table S3. Guides were cloned into the pX330 plasmid as specified in ((Cong et al. 2013); Addgene plasmid #42230). To delete EZH2, Parent ESCs were seeded at 0.5x10^6^ cells per well in a 6-well plate. The following day, the cells were transiently transfected using Lipofectamine 3000 (Invitrogen L3000-015): 800ng of sgRNA plasmid pool, or as a control, empty pX330 plasmid, and 200ng puro resistant GFP plasmid (1μg total) were mixed with 2μL P3000 reagent, 7.5μL Lipofectamine 3000 reagent and with Opti-MEM media (Gibco #31985-070) to final volume of 250μL. The reagents were incubated for 5 minutes at room temperature before being added to cells with fresh media. After 24 hours, cells were pulsed with Puromycin (2μg/mL) for 48 hours. After puromycin selection, cells were trypsinized to single cells and plated onto irradiated fibroblast feeder cells (500-2000 cells / 10cm plate) until individual colonies were visible by eye (4-5 days). Individual colonies were then selected and grown in individual wells for genotyping.

The two KO lines that were selected for further study, along with the WT control line, were then rendered dox-inducible by transfection of the rtTA-expressing plasmid described in (Kirk et al. 2018). One day prior to transfection, parent and KO ESCs were seeded at 0.5x10^6^ cells per 6-welled well. The following day, 500ng of rtTA plasmid and 500ng of transposase (1μg total DNA) were mixed with 2μL P3000 reagent, 7.5μL Lipofectamine 3000 reagent and with Opti-MEM media (Gibco #31985-070) to 250μL. The reagents incubated for 5 minutes at room temperature before being added to cells with fresh media. After 24 hours, cells underwent G418 selection (200μg/mL) for twelve days. Guides have been deposited to Addgene.

#### PCR

Genomic DNA was collected from 0.8x10^6^ cells with 500μL lysis buffer (100mM Tris-HCl pH8.1, 0.5mM EDTA pH 8.0, 200nM NaCl, 0.2% SDS) + 80μL Proteinase K (Denville) + 8μL linear acrylamide (Thermo Fisher AM9520) and incubated at 55°C overnight. Twice the volume of ice cold 100% ethanol was added. Samples were then vortexed and rotated at 4°C for 15 min. Samples were spun at max speed for 5 min at 4°C. The lysis buffer/ethanol mixture was then removed, and the DNA pellet was washed with 70% ethanol, after which the DNA pellet was resuspended in 1x TE (10mM Tris pH8.0, 1mM EDTA) and incubated overnight at 56°C. DNA concentration was measured via Nanodrop and diluted to 50ng/μL. PCR was performed with ChoiceTAQ (Denville CB4050) as follows: 25μL PCR reaction mixture (2.5μL 10x PCR reaction buffer, 0.2μL 10mM dNTPs, 0.25μL 100uM primers, 0.25μL Choice TAQ polymerase, 3μL DNA template (50ng/μL) and ddH2O to 25μL) ran in BioRad C1000 Touch or T100 thermocycler (initial denaturation at 95°C for 3 min; 25 cycles of 95°C for 30s, 60-63°C annealing for 30s, and 72°C for 30-45s extension time). PCR primers and conditions are in Table S3.

#### Total RNA isolation

ESCs were grown in 6-well plates to ∼80% confluency. Cells were washed twice with 1x PBS and 1mL of TRIzol (Thermo Fisher #15596018) was added per well. Samples were pipetted up and down at least 10 times, transferred to a microcentrifuge tube and briefly vortexed. Samples were incubated at RT for 5 minutes then 200μL of chloroform was added. Afterwards, samples were vortexed and incubated for 3 minutes at RT. Samples were spun down at 12,000 rpm for 15 min at 4°C. The upper aqueous phase was moved to a new tube and 8μL of linear acrylamide (Thermo Fisher, AM9520) was added. Then 500μL of 100% isopropanol was added, and samples were vortexed and incubated at RT for 10 minutes. Tubes were spun down 12,000 rpm for 10 minutes at 4°C. Supernatant was removed and pellets washed with 1mL of cold 75% ethanol. Samples were briefly vortexed and spun down at 7,500 rpm for 5 min at 4°C. Supernatant was discarded and pellet was dried by repeated spin down and aspiration. Final pellets were resuspended in 100μL water.

#### RT-qPCR

Equal amounts of RNA (0.5–1μg) were reverse transcribed using the High-Capacity cDNA Reverse Transcription Kit (Thermo Fisher Scientific, #4368813) with the random primers provided, and then diluted with 30μL 1xTE. For RIP RT-qPCR, 2μL of eluted sample were used in RT reactions. 10μL qPCR reactions were performed using iTaq Universal SYBR Green (Bio-Rad) and custom primers on a Bio-Rad CFX96 system with the following thermocycling parameters: initial denaturation at 95°C for 10 min; 40 cycles of 95°C for 15s, 60°C for 30s, and 72°C for 30s followed by a plate read. The primer concentration used for all qPCR reactions in this study was 0.5μM. Standard curves were used in all qPCR analyses and were prepared by RT of equal volume of wild type sample to other samples. After RT, five 5-fold serial dilutions were made (6 total standards including undiluted RT reaction) and added in duplicate to qPCR plates. After qPCR run, samples were normalized to standard curve read using the BioRad CFX Manager Software. See Table S3 for all primer sequences used.

#### Antibodies

All antibodies and their conditions used for this study are listed in Table S4.

#### Formaldehyde crosslinking of ESCs

For RIP, cells were grown to 75-85% confluency, trypsinized and counted. Cells were washed twice in cold 1xPBS then rotated for 30 min in 10mL of 0.3% formaldehyde (1mL 16% methanol-free formaldehyde (Pierce, #28906) in 49mL 1xPBS) at 4°C. Formaldehyde was quenched with 1mL of 2M glycine for 5 min at room temperature with rotation. Cells were washed 3x in cold 1xPBS, then resuspended in 1xPBS at 10x10^6^ cells per mL, aliquoted 10x10^6^ cells to new tubes, and spun down. PBS was aspirated and pellets were snap frozen in a liquid nitrogen bath and immediately transferred to -80°C until use.

#### Formaldehyde + DSG crosslinking of ESCs for EZH2-CS5246 ChIP

Cells were grown to 80% confluency and washed three times in room temperature 1xPBS. Cells were scraped and transferred to conical tubes, spun down and supernatant removed. Cells were first crosslinked with 15 mL of 2mM DSG (Thermo Fisher #20593) in 1xPBS and rotated at RT for 45 min. Cells were then washed two times in room temperature 1xPBS and then crosslinked for 15 minutes at room temperature with 15mL of 1% formaldehyde (Thermo Fisher #28906) with rotation. Formaldehyde was quenched with 1.67mL of 2.5M glycine for five minutes with rotation. Conical tubes were placed on ice and cells were washed twice with cold 1xPBS and then spun down at 3,000 rpm at 4°C. Cell pellets were resuspended in 1xPBS supplemented with 1mM PMSF and 1x Protease Inhibitor Cocktail to a concentration of 10x10^6^ cells/mL. Cells were aliquoted 10x10^6^ cells to new tubes. PBS was spun out at 1,200 rpm for 5 mins at 4°C. PBS was aspirated, and cells were flash frozen with a dry ice-methanol bath and stored at -80°C until use.

#### RNA-IPs (RIPs)

RIPs were performed similar to (Schertzer et al. 2019a), which is a protocol originally adapted from (Hendrickson et al. 2016; Raab et al. 2019). 25μL protein A/G agarose beads (Santa Cruz sc-2003) were washed three times in blocking buffer (0.5% BSA in 1xPBS) and incubated overnight at 4°C with 10μL antibody (anti-SAFB, Bethyl 812-300A; EZH2, Cell Signaling #5246) or 10μg IgG (Invitrogen #02-6502). 10x10^6^ cells were resuspended in 500μL RIPA Buffer (50mM Tris-HCl, pH8, 1% Triton X-100, 0.5% sodium deoxycholate, 0.1% SDS, 5mM EDTA, 150mM KCl) supplemented with 1x Protease Inhibitor Cocktail (PIC; Sigma #P8340), 2.5μL SuperaseIN (Thermo Fisher AM2696) and 0.5mM DTT (ThermoFisher #15508013) and sonicated twice for 30s on and 1 min off at 30% output using the Sonics Vibracell Sonicator (Model VCX130, Serial# 52223R). Samples were spun down at high speed for 15 minutes and 50μL total lysate was saved for input. Beads were washed three times in 1mL fRIP buffer (25mM Tris-HCl pH7.5, 5mM EDTA, 0.5% NP-40, 150mM KCl) and resuspended in 450μL fRIP buffer supplemented as above with PIC, SuperaseIN and DTT, then mixed with sonicated samples. Samples were rotated overnight at 4°C, then washed once with 1mL fRIP buffer and resuspended in 1mL PolII ChIP Buffer (50mM Tris-HCl pH 7.5, 140mM NaCl, 1mM EDTA, 1mM EGTA, 1% Triton X-100, 0.1% Sodium-deoxycholate, 0.1% Sodium dodecyl sulfide) before being transferred to a new tube. Samples were rotated at 4°C for five minutes, spun down at 1,200 rpm, and the supernatant aspirated. Samples were washed twice more with 1mL PolII ChIP Buffer, once with 1mL High Salt ChIP Buffer (50mM Tris-HCl pH7.5, 500mM NaCl, 1mM EDTA, 1mM EGTA, 0.1% sodium-deoxycholate, 0.1% sodium dodecyl sulfide, 1% Triton X-100), and once in 1mL LiCl buffer (20mM Tris pH8.0, 1mM EDTA, 250mM LiCl, 0.5% NP-40, 0.5% sodium-deoxycholate); each wash included a five-minute rotation at 4°C. At the final wash, samples were transferred to a new tube. After the final wash, inputs were thawed on ice and bead samples were resuspended in 56μL water, 33μL of 3x reverse-crosslinking buffer (3x PBS, 6% N-lauroyl sarcosine and 30mM EDTA), 5μL 100mM DTT (ThermoFisher #15508013), 20μL Proteinase K (Thermo Fisher #25530015), and 1μL of SuperaseIN. Samples were incubated for 1hr at 42°C, 1hr at 55°C, 30 minutes at 65°C, and mixed by pipetting every 15 minutes. Afterwards, 1mL Trizol (Thermo Fisher #15596018) was added, samples were vortexed, 200μL chloroform was added, samples were vortexed again, and finally spun at 12,000 rpm for 15 mins at 4°C. The aqueous phase was then extracted and to that one volume of 100% ethanol was added. Samples were vortexed and applied to Zymo-Spin IC Columns (Zymo #R1014) and spun for 30 seconds at top speed on a benchtop microcentrifuge. 400μL of RNA Wash Buffer (Zymo #R1014) was added and samples were spun at top speed for 30 seconds. For each sample, 5μL DNase I (Zymo #R1014) and 35μL of DNA Digestion Buffer (Zymo #R1014) was added directly to the column matrix and incubated at room temp for 20 minutes. 400μL of RNA Prep Buffer (Zymo #R1014) was then added to the columns, and columns were spun at top speed for 30 seconds. 700μL RNA Wash Buffer (Zymo #R1014) was then added, and columns were spun at top speed for 30 seconds. 400μL RNA Wash Buffer was then added, and columns were spun at top speed for 30 seconds. The flow through was discarded and columns spun again for 2 minutes to remove all traces of wash buffer. Columns were transferred to a clean tube, 15μL of ddH2O was added to each column, and after a five-minute incubation, samples were spun at top speed for two minutes to elute.

#### RNA sequencing

For RIP-Seq samples, 9μL of RIP sample (from 15μL total) were used. Each library preparation included 1μL of 1:250 dilution of ERCC Spike-In RNAs (Ambion #4456653). 10μL total were prepped using the KAPA RNA HyperPrep Kit with RiboErase (Kapa Biosystems; product #KR1351). Sequencing was performed on an Illumina Next-Seq 500, using high-output, 75-cycle kits (Illumina #20024906).

#### EZH2 ChIP-Seq

The day before sonication, 25μL of protein A/G agarose beads (Santa Cruz sc-2003) were washed 3 times in block solution (0.5% BSA in 1xPBS) before being resuspended in 300μL blocking solution. 10μL EZH2 antibody (Cell Signaling #5246) per 10 million cells was added, then beads and antibody conjugated via rotation overnight at 4°C. On the day of sonication, 10 million ESCs crosslinked with DSG and 1% formaldehyde were resuspended in PolII ChIP Buffer (no Triton) (50mM Tris-HCl, pH 7.5, 140mM NaCl, 1mM EDTA, pH 8.0, 1mM EGTA, pH 8., 0.1% Sodium-deoxycholate, 0.1% Sodium dodecyl sulfide) supplemented with 1x Protease Inhibitor Cocktail (PIC; Sigma Product #P8340) and then sonicated ten times for 30s on and 1 min off at 30% output using the Sonics Vibracell Sonicator (Model VCX130, Serial# 52223R). Supernatant was cleared by top speed centrifugation for 20 minutes, moved to a new tube and 10% Triton-X100 added to a final 1%. 10% of final volume was taken and saved as input. Conjugated antibody/bead mixture was spun down and washed three times in blocking buffer. ChIPs were then performed by incubating sonicated cell lysates with pre-conjugated EZH2 antibody/agarose beads overnight at 4°C. The next day, samples were spun down and resuspended 1mL PolII ChIP Buffer (50mM Tris-HCl pH 7.5, 140mM NaCl, 1mM EDTA, 1mM EGTA, 1% Triton X-100, 0.1% Sodium-deoxycholate, 0.1% Sodium dodecyl sulfide) before being transferred to a new tube. Samples were rotated at 4°C for five minutes, spun down at 2,000 rpm for two minutes, and the supernatant aspirated. Samples were washed twice more with 1mL PolII ChIP Buffer, once with 1mL High Salt ChIP Buffer (50mM Tris-HCl pH 7.5, 500mM NaCl, 1mM EDTA, 1mM EGTA, 0.1% sodium-deoxycholate, 0.1% sodium dodecyl sulfide, 1% Triton X-100), once in 1mL LiCl buffer (20mM Tris pH 8.0, 1mM EDTA, 250mM LiCl, 0.5% NP-40, 0.5% sodium-deoxycholate), and once in 1xTE; each wash included a five-minute rotation at 4°C. At the final wash, samples were transferred to a new tube. To elute the DNA, beads were re-suspended in 250μL Elution buffer (50mM Tris pH 8.0, 10mM EDTA, and 1% SDS) and placed on a 65°C heat block for 17 minutes with vortexing every 2-3 minutes. Supernatant was moved to a new tube, 240μL and 10μL 5M NaCl, and crosslinks were reversed overnight at 65°C. The following day, eluates were incubated with 3μL RNase A (Thermo Fisher EN0531) for one hour at 37°C and then 10μL Proteinase K (Thermo Fisher #25530015) for two to three hours at 56°C. One volume (510μL) of phenol:chloroform:isoamyl alcohol (Sigma P3803) was added to the DNA, and was spun at max speed for 5 mins at 4°C. The aqueous solution was transferred to a new tube. 5μL of linear acrylamide was added, and then 1/10 volume (50μL) of 3M NaOAc, pH5.4. Samples were vortexed and spun down. 2 volumes (1mL) of ice cold 100% Ethanol was added, and samples were vortexed, spun down and incubated overnight at -20°C. The next day, samples were spun at max speed for 30 mins at 4°C (cold room). Supernatant was aspirated and DNA pellet was washed in 1mL ice-cold 80% Ethanol. Samples were spun down and supernatant was removed. Samples were dried by repeated aspiration and pulse spin. Samples were resuspended in 30μL 1X TE for sequencing. DNA was prepared for sequencing on the Illumina platform using Next Reagents (NEB) and Agencourt AMPure XP beads (Beckman Coulter).

### Computational analyses

#### Mass spectrometry data analysis

Raw data files were processed using MaxQuant version 1.6.12.0 and searched against the reviewed mouse database (containing 17,051 entries), appended with a contaminants database, using Andromeda within MaxQuant. Enzyme specificity was set to trypsin, up to two missed cleavage sites were allowed, and methionine oxidation and N-terminus acetylation were set as variable modifications. A 1% FDR was used to filter all data. Match between runs was enabled (5 min match time window, 20 min alignment window), and a minimum of two unique peptides was required for label-free quantitation using the LFQ intensities. Perseus was used for further processing (Tyanova et al. 2016). Only proteins with >1 unique+razor peptide were used for LFQ analysis. Proteins with 50% missing values were removed and missing values were imputed from normal distribution within Perseus. Log2 fold change (FC) ratios were calculated using the averaged Log2 LFQ intensities of IP sample compared to IgG control, and students t-test performed for each pairwise comparison, with p-values calculated. Proteins with significant p-values (<0.05) and Log2 FC >1 were analyzed further. All analyzed protein interaction data are present in Table S1. The mass spectrometry proteomics data have been deposited to the ProteomeXchange Consortium via the PRIDE partner repository (Perez-Riverol et al. 2022) with the dataset identifier PXD038103. Reviewer account details: [Username; Password: 7TiboSU8].

#### RIP-Seq alignment

RIP-Seq data were aligned to the mm10 mouse genome using STAR with default parameters (Dobin et al. 2013). Alignments with a quality score ≥30 were retained for subsequent analyses (Li et al. 2009).

#### RIP-Seq peak calling

SAFB peaks were identified as part of (Cherney et al. 2022). For EZH2 RIP-Seq peak calling, after alignment and filtering, all EZH2 RIP-Seq data from WT ESC replicates were concatenated and using samtools, split into two files, corresponding to alignments that mapped to the positive and negative strands of the genome, respectively. Using a custom perl script, the strand information within the positive and negative strand alignment files was randomized so as to better match the criteria of the MACS peak caller, which uses the average distance between positive and negative strand alignments to estimate the fragment length (Zhang et al. 2008). Putative peaks were called on strand-randomized positive and negative strand alignment files, respectively, using default MACS parameters and not providing a background file (Zhang et al. 2008). Peak bed files were converted to SAF format and reads under each putative peak were counted from EZH2 RIP-Seq alignments performed in WT, EZH2 KO, and SAFB/2 DKO ESCs using featureCounts (Liao et al. 2014). EZH2 peaks were determined by retaining putative peaks that were represented by at least five reads in at least two of the three WT ESC lines profiled. We retained only those putative peaks that harbored an average aligned-reads-per-million-total-reads (RPM) signal of at least five-fold less in IgG RIPs compared to wild-type EZH2 RIPs. This yielded 33,671 regions that were enriched in their association with EZH2 in wild-type ESCs. (Table S2).

#### Significance of RIP-Seq peak overlaps

To determine the significance of RIP-Seq peak overlaps, the hypergeometric test in R (Team 2021) was used under the following conditions: phyper(q=overlap, m=set1 (post-filter), n=total peaks (pre-filter) – set1 (post-filter), k=set2 (post-filter), lower.tail = FALSE).

#### RIP-Seq scatter plots

Scatter plots in Figure 2 were constructed using featureCounts to count the reads under each of the 30,998 SAFB peaks in each dataset (Liao et al. 2014). Read counts were then plotted using R (Team 2021).

#### UCSC wiggle density plots

UCSC wiggle density plots were made from filtered sam files using custom perl scripts normalized to reads per million (rpm). Tracks of pooled replicates are located here: https://genome.ucsc.edu/s/recherney/Cherney_Antibody_2023

#### ChIP-Seq peak calling

EZH2 data were aligned to mm10 using bowtie2 (Langmead and Salzberg 2012). Peaks were called using MACS2 under the following parameters using an H3 control: [macs2 callpeak -t -c bam -n -f BAM -g mm –broad –broad-cutoff 0.01] (Zhang et al. 2008). EZH2 peak locations are included in Table S5.

#### ChIP-Seq scatter plots

Scatter plots in Figure 4 were constructed using featureCounts to count the reads from each EZH2 ChIP-seq data set under each of the EZH2 Cherney peaks (Table S5; (Liao et al. 2014)). Read counts were then plotted using R (Team 2021).

#### Sequencing data

https://www.ncbi.nlm.nih.gov/geo/query/acc.cgi?acc=GSE227893 – ChIP-Seq and RIP-Seq Secure token for GSE227893: ilobcwuapponbwz

## Funding

This work was supported by NIH National Institute of General Medical Sciences (NIGMS) grant number R01GM136819 to J.M.C, T32 GM007092 and Eunice Kennedy Shriver National Institute of Child Health and Human Development (NICHD) grant number F31 HD103334 to R.E.C, and T32 GM119999 and NICHD grant number F31 HD103370 to A.K.B. The proteomics work was conducted at the UNC Proteomics Core Facility, which is supported in part by NCI Center Core Support Grant (2P30CA016086-45) to the UNC Lineberger Comprehensive Cancer Center.

## Acknowledgements

The authors would like to thank UNC colleagues for many helpful discussions.

## Competing Interests

The authors declare no competing interests.

## Notes

### Competing Interest Statement

The authors have declared no competing interest.

